# Reliability Assessment of Tissue Classification Algorithms for Multi-Center and Multi-Scanner Data

**DOI:** 10.1101/2020.01.28.922971

**Authors:** Mahsa Dadar, Simon Duchesne, For the CCNA Group and the CIMA-Q Group

**Author notes:** **Corresponding Author Information:** Mahsa Dadar, Cervo Brain Research Centre, 2601 Chemin de la Canardière, Québec, Canada, G1J 2G3.

## Abstract

**Background:** Gray and white matter volume difference and change are important imaging markers of pathology and disease progression in neurology and psychiatry. Such measures are usually estimated from tissue segmentation maps produced by publicly available image processing pipelines. However, the reliability of the produced segmentations when using multi-center and multi-scanner data remains understudied. Here, we assess the robustness of six publicly available tissue classification pipelines across images acquired from different MR scanners and sites.

**Methods:** We used T1-weighted images of a single individual, scanned in 73 sessions across 27 different sites to assess the robustness of the tissue classification tools. Variability in Dice Kappa values and tissue volumes was assessed for Atropos, BISON, Classify_Clean, FAST, FreeSurfer, and SPM12. We also estimated the sample size necessary to detect a significant 1% volume reduction based on the variability of the estimates from each method within and across scanner models.

**Results:** BISON had the lowest overall variability in its volumetric estimates, followed by FreeSurfer, and SPM12. All methods also had significant differences between some of their estimates across different scanner manufacturers (e.g. BISON had significantly higher GM estimates and correspondingly lower WM estimates for GE scans compared to Philips and SIEMENs), and different signal-to-noise ratio (SNR) levels (e.g. FAST and FreeSurfer had significantly higher WM volume estimates for high versus medium and low SNR tertiles as well as correspondingly lower GM volume estimates). BISON also had the smallest sample size requirement across all scanners and tissue types, followed by FreeSurfer, and SPM12.

**Conclusions:** Our comparisons provide a benchmark on the reliability of the publicly used tissue classification techniques and the amount of variability that can be expected when using large multi-center datasets and multi-scanner databases.

## 1. Introduction

Multi-center studies, usually performed to increase sample sizes, provide researchers with a plethora of data to explore different hypotheses with sufficient statistical power. However, such datasets bring about their own new set of challenges, particularly when acquired without harmonization and on different scanner platforms (Duchesne et al., 2019a). In particular, the dynamic range and intensity characteristics of the images produced by different scanner models, from different vendors and operated in different configurations, might significantly vary across acquisition sites. Such differences can in turn impact the reliability of tissue classification, one of the most commonly performed tasks in structural neuroimaging studies (e.g. for purposes of voxel based morphometry) or as a necessary step in other post-processing pipelines (e.g. for purposes of cortical thickness extraction or diffusion tensor imaging) (González-Villà et al., 2016; Mateos-Pérez et al., 2018). Intensity variations may lead to systematic biases in tissue classification when using or comparing data across different centers, and hence adversely influence the final results.

There have been demonstrations of the potential impact of scanner variability on estimates for tissue classification in neuroimaging pipelines. Using data from two travelling human phantoms across four different sites, Gouttard et al. assessed the variability in tissue classification and voxel based morphometry across sites, reporting high intra-scanner variabilities, as well as higher inter-scanner variabilities, between Siemens Healthcare’s Allegra and Tim Trio scanners (Gouttard et al., 2008). Similarly, using data collected on four different scanner models at five different sites in six different subjects, Schnak et al. showed a significant site effect in gray and white matter segmentations, voxel-based morphometry, and cortical thickness measurements (Schnack et al., 2004, 2010). Similarly, Pardoe et al. reported significant site-specific differences in voxel-based morphometry measurements between healthy control subjects scanned across three different sites (Pardoe et al., 2008).

However, no studies have comprehensively compared the performance of commonly used tissue classification methods on different scanner models from three of the most commonly used platforms in clinical and research settings (i.e. GE Healthcare (MI, USA), Siemens Healthcare (Erlangen, GER), and Philips Medical Systems (Best, NED)). Using data from the Single Individual volunteer for Multiple Observations across Networks (SIMON) public dataset, we had a unique opportunity to perform such comparison across 90 scans of this single individual, acquired within the span of seven years on 28 different sites with 12 different scanner models. In this study, we compared the performance of six publicly available, widely used tissue classification methods (Atropos (Avants et al., 2011); BISON (Dadar and Collins, 2019); Classify_Clean (Cocosco et al., 2003); FAST (Zhang et al., 2001); FreeSurfer (Fischl, 2012); and SPM12 (Ashburner et al., 2014, p. 12). These comparisons provide a benchmark on the reliability of each technique, and the amount of variability that can be expected when using multi-center datasets. We hypothesized that (a) there would be statistically significant differences between tissue volumes estimated by the methods across scanner manufactures; and (b) the signal to noise ratio (SNR) level of the images would have a statistically significant effect on estimated volumes. Finally, we estimated the sample size necessary to detect a 1% reduction in tissue volumes with sufficient power based on each method within and across scanner models.

## 2. Methods

### 2.1. Data

Data used in this study included 90 3T T1-weighted MRIs from the SIMON dataset, a sample of convenience of one healthy male aged between 39 and 46 years old, scanned for research projects in 73 sessions at 28 sites on a variety of platforms, namely: GE Healthcare (DISCOVERY MR750 and SIGNA Pioneer); Philips Medical Systems (Achieva, Ingenia, Intera, and T5); and Siemens Healthcare (Allegra, Prisma, PrismaFit, Skyra, SonataVision, Symphony, and TrioTim) (Duchesne et al., 2019a, 2019b). The data was acquired with a number of different protocols, but more than two-thirds complied with the harmonized Canadian Dementia Imaging Protocol (www.cdip-pcid.ca; (Duchesne et al., 2019a)). For more information and access to the dataset, see (http://fcon_1000.projects.nitrc.org/indi/retro/SIMON.html).

### 2.2. Image Processing

All T1-weighted scans were processed through standard preprocessing steps using the MINC toolkit (https://github.com/BIC-MNI/minc-tools): denoising (Coupe et al., 2008), intensity non-uniformity correction (Sled et al., 1998), and intensity normalization into range (0-100). All images were then linearly registered to MNI-ICBM152 template at an isotropic 1×1×1 mm^3^ resolution (Collins and Evans, 1997; Dadar et al., 2018) to enable comparisons between segmented tissue masks. Nonlinear registration to the MNI-ICBM152 template was also performed at 1×1×1 mm^3^ resolution using ANTs (Avants et al., 2008). Brain extraction was performed on the linearly registered images using BEaST (Eskildsen et al., 2012). To avoid any variability in the estimates caused by potential differences between BEaST masks obtained from different scans, a single brain mask was generated by intersecting the BEaST brain masks for all scans. All segmentations were consistently compared inside this single brain mask. To investigate the effect of signal to noise ratio (SNR) on the segmentations, SNR from each image was obtained using a robust Rician noise estimation technique (Coupé et al., 2010).

### 2.3. Tissue Segmentation

Tissue segmentations were performed after these preprocessing steps using: 1) Atropos (Avants et al., 2011); 2) BISON (Dadar and Collins, 2019); 3) Classify_Clean (Cocosco et al., 2003); FAST 5.0 (Zhang et al., 2001); FreeSurfer 6.0.0 (Fischl, 2012); and SPM12 (Penny et al., 2011). For all pipelines, default settings were used.

#### 2.3.1. ANTs Atropos

Atropos is an open-source multi-class segmentation pipeline which performs tissue classification using a Bayesian framework, incorporating template-based tissue probability maps in the form of Markov Random Fields (Avants et al., 2011). Atropos is publicly available at https://github.com/ANTsX/ANTs/blob/master/Scripts/antsAtroposN4.sh. Tissue probability maps for Atropos were generated by registering MNI-ICBM152 tissue priors to the subject’s brain using the estimated nonlinear registrations. Atropos was then run for 3-dimension inputs with 3 classes using the following command and parameters:

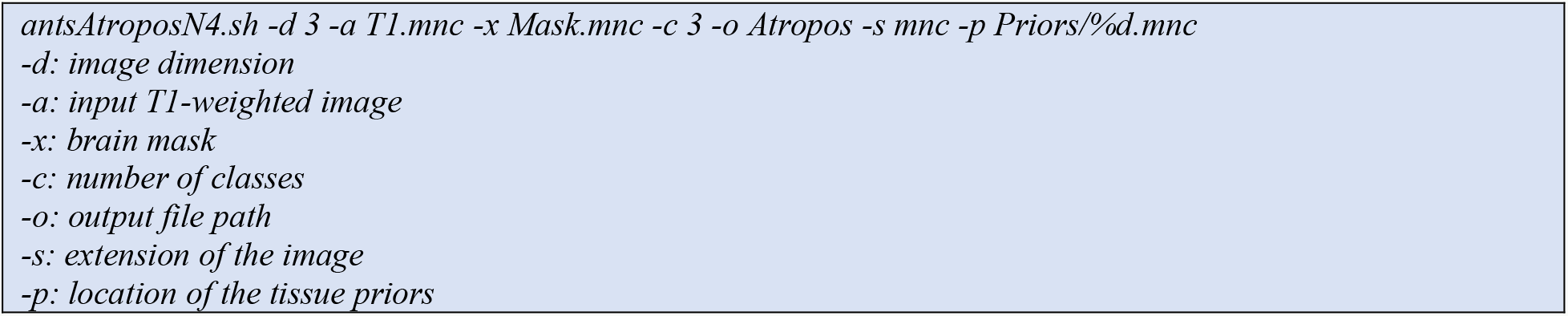

#### 2.3.2. BISON

Brain tISue segmentatiON (BISON) is an open source pipeline based on a random forests classifier that has been trained using a set of intensity and location features from a multi-center manually labelled dataset of 72 individuals aged from 5-96 years (Dadar and Collins, 2019). The BISON script as well as a pretrained random forest classifier is publicly available at http://nist.mni.mcgill.ca/?p=2148. BISON was run using the following command:

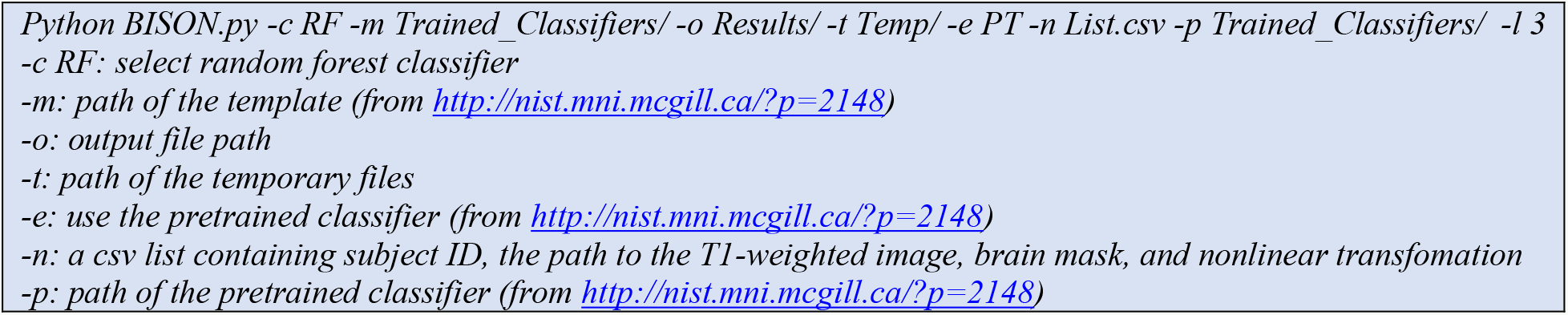

#### 2.3.3. Classify_Clean

Classify_Clean (Cocosco et al., 2003) is an executable provided as part of the MINC toolkit. It uses a set of standard sample points to compute an initial classification, which is then used to purge incorrect tag points. The resulting tag point set is used by a neural network classifier to perform tissue segmentation. Classify_Clean was run using the following command:

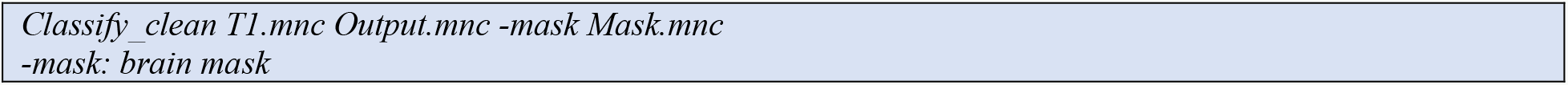

#### 2.3.4. FAST

FMRIB’s Automated Segmentation Tool (FAST) performs tissue classification while also correcting for intensity inhomogeneity (Zhang et al., 2001). FAST is based on a hidden Markov Random Field model and an associated Expectation-Maximization algorithm. To achieve optimal results, the T1-weighted images were first masked. FAST 5.0 was then run using the following command:

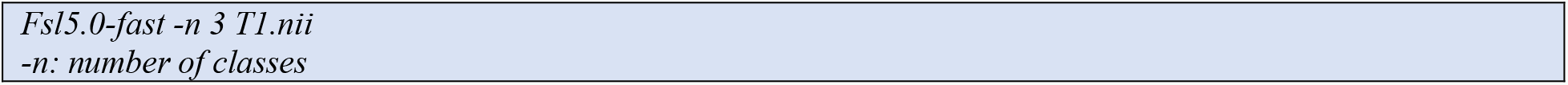

#### 2.3.5. FreeSurfer

FreeSurfer is an open source brain image processing software that provides a full processing stream for structural T1-weighted data (Fischl, 2012). FreeSurfer is publicly available at https://surfer.nmr.mgh.harvard.edu/. Since FreeSurfer performs its own preprocessing, to achieve optimal results, unpreprocessed T1-weighted images were run by FreeSurfer, and the final segmentation output (aseg.mgz) was then used to generate a tissue classification map using the FreeSurfer Look Up Table of the segmented regions available at https://surfer.nmr.mgh.harvard.edu/fswiki/FsTutorial/AnatomicalROI/FreeSurferColorLUT. FreeSurfer 6.0.0 was run using the following command:

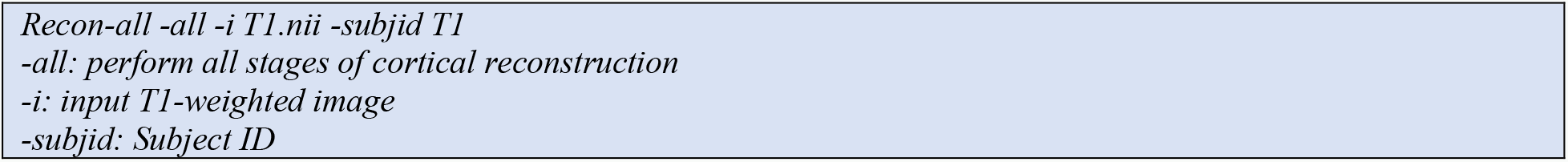

#### 2.3.6. SPM12

Statistical Parametric Mapping (SPM) performs tissue segmentation by estimating the nonlinear deformation field that overlays tissue probability maps to new image (Ashburner et al., 2014, p. 12). After this step, a few iterations of a simple Markov Random Field procedure are run to clean up the final results. To generate a single tissue classification mask from the resulting probabilistic output maps for GM (c1T1.nii), WM (c2T1.nii), and CSF (c3T1.nii), the label with the highest probability from the three classes was assigned to each voxel. Tissue segmentation was performed using the SPM12 GUI, with default parameters.

### 2.4. Comparisons and Statistical Analyses

To generate a silver standard segmentation as a benchmark for comparison, an average T1-weighted image template (the “template”) was created out of all original scans using a previously validated method for generating unbiased average templates (Figure S.1) (Fonov et al., 2009, 2011). For each algorithm, we then generated the segmentation mask for this method on the template, as a silver standard against which to compare other segmentations. Dice Kappa similarity index (Dice, 1945) was used to compare the segmentations against the silver standard. To assess the statistical significance of the results, paired *t*-tests were performed on the Dice Kappa values of all pairs of segmentation techniques, and the resulting p-values were corrected for multiple comparisons using false discovery rate (FDR).

Volumetric comparisons were also used to assess the performance of the six methods. Prior to the analyses, all tissue volumes were normalized with respect to the total intracranial volume (ICV), estimated based on the single brain mask (BEAST; section 2.2). To assess whether scanner differences and SNR systematically affect the segmentations produced by each method, the following linear regression models were tested:

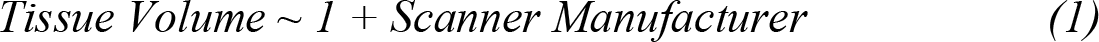

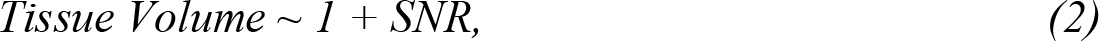

where Scanner Manufacturer is a categorical variable reflecting GE, Siemens, and Philips manufacturers, and SNR is a categorical variable reflecting low, medium, and high SNR (instead of using SNR as a continuous variable, categorical SNR tertitles were used due to the non-normal distribution of the SNR values). All results were corrected for multiple comparisons using FDR with a significance threshold of 0.05.

The sample sizes (per arm) necessary to detect a 1% reduction in the tissue volumes were estimated using the sampsizepwr function from MATLAB (80% power, 2-tailed significance, p = 0.05). For the within scanner analyses, the standard deviations were adjusted for the sample size:

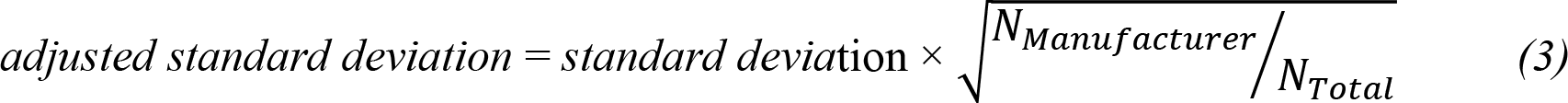

where *N*_*Manufacturer*_ denotes the number of scans from a specific manufacturer and *N*_*Total*_ denotes the total number of scans. All analyses were performed using MATLAB version 2019b.

## 3. Results

Figure 1 shows the axial slices of the 90 scans after preprocessing, linear registration to the MNI-ICBM152 template, and brain extraction.

**Figure 1.**
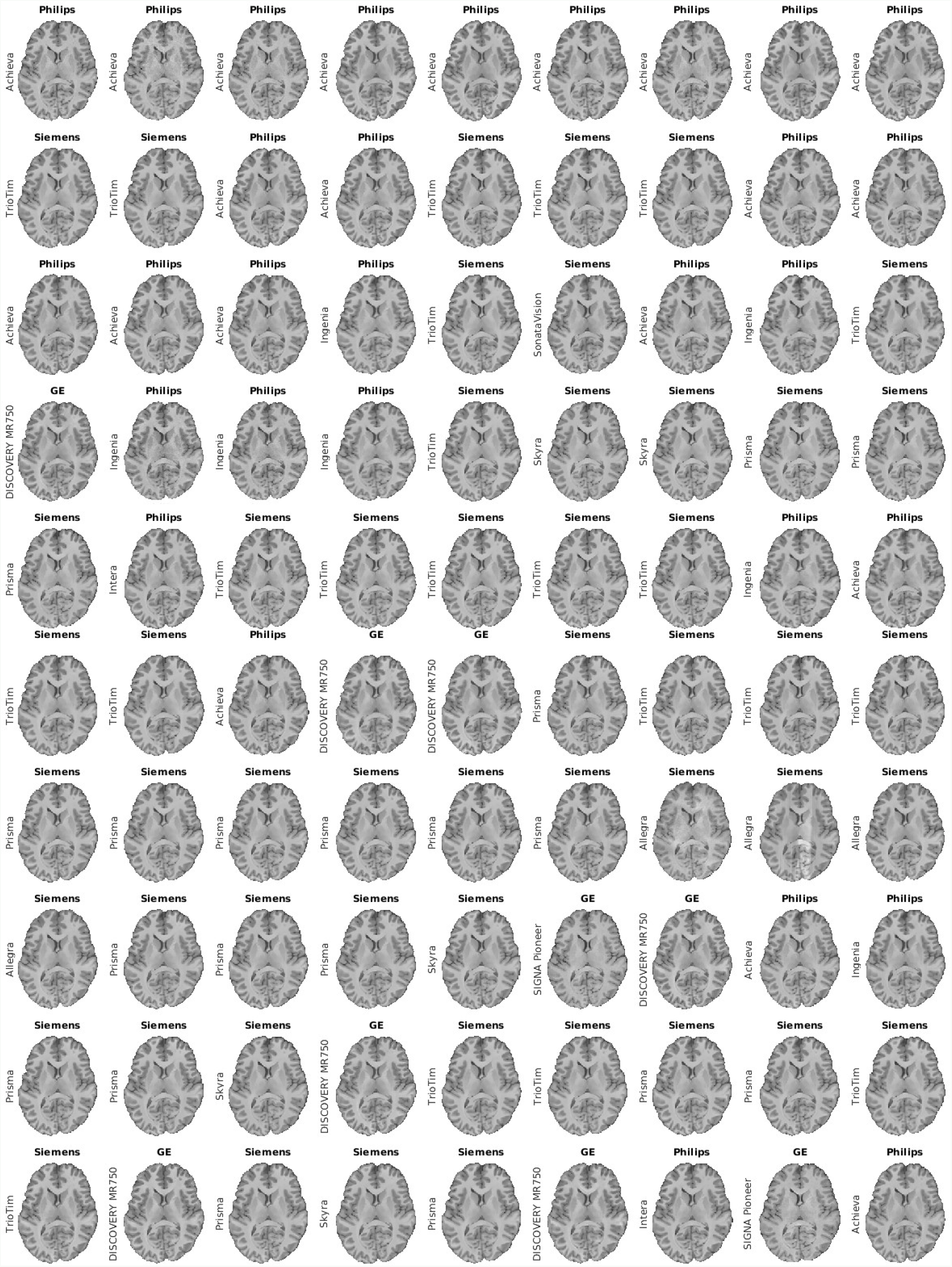
Axial slices of the preprocessed images, after denoising, inhomogeneity correction, intensity normalization, and brain extraction. Scans are ordered chronologically.

FAST failed to produce an output for two of the images, while FreeSurfer did not produce an output for one case with some imaging artefacts (Figure 1, row 7, column 8). The latter case was removed from all analyses. All other tools processed all images successfully.

Figure 2 shows axial slices of the average template as well as the silver standard tissue segmentation masks generated by each method. Figure S.2 shows axial slices covering the brain overlayed with segmentations from each method, for one scan example (Philips Intera scanner 3T). Table 1 shows the average overall Dice Kappa comparing each segmentation against the silver standard as well as the average Dice Kappas separately for each manufacturer. Figure 3 shows boxplots of the Dice Kappas for each tissue type across scanner manufacturers.

**Figure 2.**
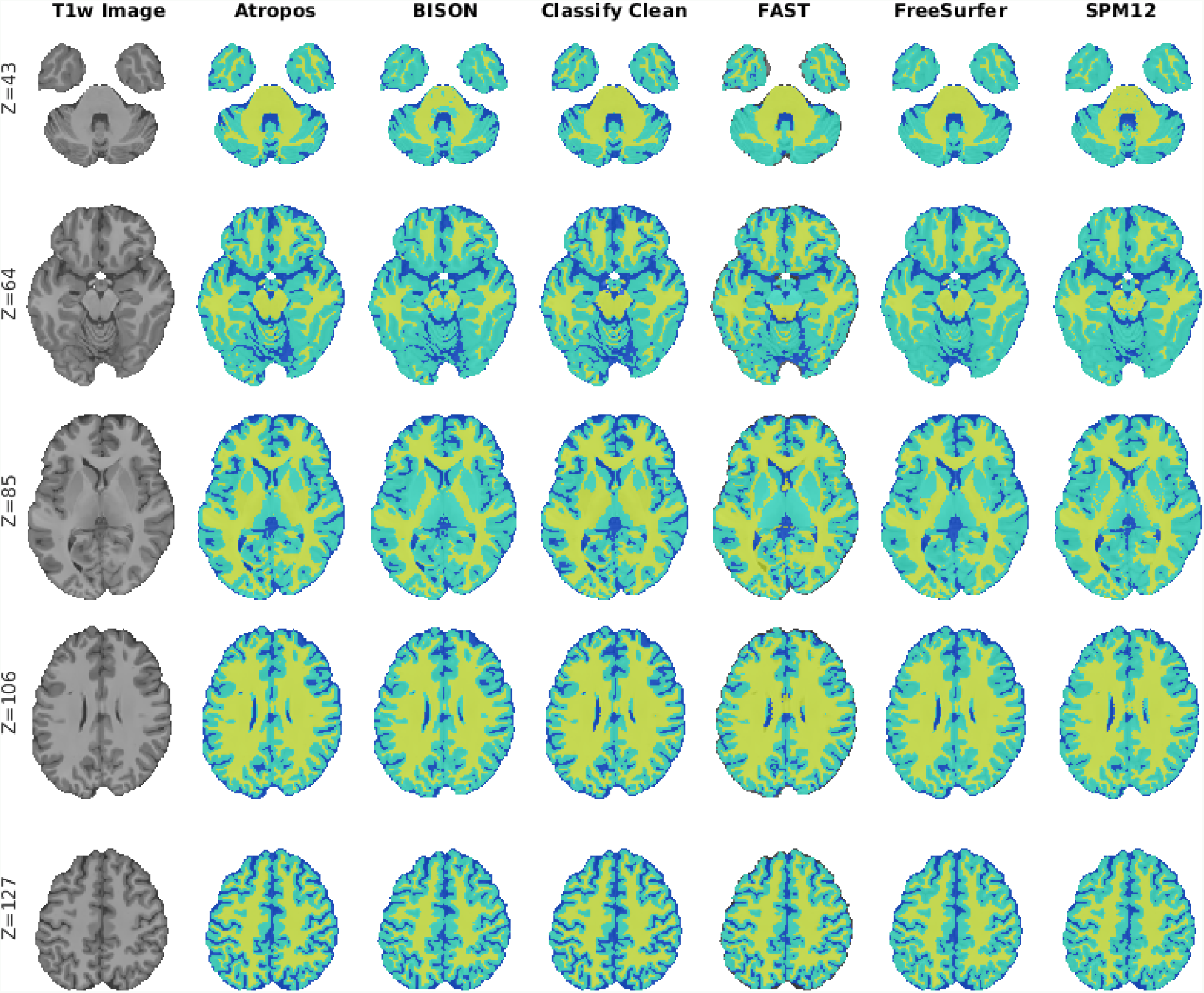
Axial slices showing the template created from the original T1w MRIs, and the silver standard segmentations of this template from Atropos, BISON, Classify_Clean, FAST, FreeSurfer, and SPM12.

**Table 1.** Average Dice Kappas for each method for all individual scans, when compared to the silver standard segmentation on the template. Values indicate mean ± standard deviation. All values are in % of total intracranial volume.

| Tissue | Gray Matter |  |  |  | White Matter |  |  |  | Cerebrospinal Fluid |  |  |  |
| --- | --- | --- | --- | --- | --- | --- | --- | --- | --- | --- | --- | --- |
| Method | Overall | GE | Philips | SIEMENS | Overall | GE | Philips | SIEMENS | Overall | GE | Philips | SIEMENS |
| Atropos | $0.92 \pm 0.03$ | $0.88 \pm 0.06$ | $0.91 \pm 0.03$ | $0.93 \pm 0.02$ | $0.94 \pm 0.02$ | $0.92 \pm 0.04$ | $0.94 \pm 0.01$ | $0.95 \pm 0.01$ | $0.85 \pm 0.03$ | $0.82 \pm 0.02$ | $0.83 \pm 0.03$ | $0.87 \pm 0.03$ |
| BISON | $0.94 \pm 0.01$ | $0.93 \pm 0.01$ | $0.93 \pm 0.01$ | $0.94 \pm 0.01$ | $0.93 \pm 0.01$ | $0.92 \pm 0.01$ | $0.93 \pm 0.01$ | $0.94 \pm 0.01$ | $0.82 \pm 0.03$ | $0.80 \pm 0.02$ | $0.81 \pm 0.02$ | $0.83 \pm 0.03$ |
| Classify | $0.91 \pm 0.02$ | $0.91 \pm 0.02$ | $0.91 \pm 0.02$ | $0.92 \pm 0.02$ | $0.92 \pm 0.02$ | $0.91 \pm 0.03$ | $0.91 \pm 0.01$ | $0.93 \pm 0.02$ | $0.81 \pm 0.03$ | $0.80 \pm 0.03$ | $0.80 \pm 0.03$ | $0.81 \pm 0.03$ |
| FAST | $0.87 \pm 0.02$ | $0.85 \pm 0.03$ | $0.87 \pm 0.02$ | $0.88 \pm 0.01$ | $0.91 \pm 0.01$ | $0.91 \pm 0.01$ | $0.91 \pm 0.01$ | $0.90 \pm 0.01$ | $0.82 \pm 0.04$ | $0.78 \pm 0.04$ | $0.80 \pm 0.04$ | $0.85 \pm 0.03$ |
| FreeSurfer | $0.84 \pm 0.01$ | $0.84 \pm 0.01$ | $0.84 \pm 0.02$ | $0.85 \pm 0.01$ | $0.89 \pm 0.01$ | $0.89 \pm 0.01$ | $0.89 \pm 0.01$ | $0.89 \pm 0.01$ | $0.69 \pm 0.03$ | $0.67 \pm 0.03$ | $0.69 \pm 0.02$ | $0.69 \pm 0.03$ |
| SPM12 | $0.93 \pm 0.01$ | $0.92 \pm 0.02$ | $0.93 \pm 0.01$ | $0.94 \pm 0.01$ | $0.93 \pm 0.01$ | $0.92 \pm 0.02$ | $0.93 \pm 0.01$ | $0.94 \pm 0.01$ | $0.84 \pm 0.04$ | $0.82 \pm 0.02$ | $0.82 \pm 0.03$ | $0.85 \pm 0.03$ |

**Figure 3.**
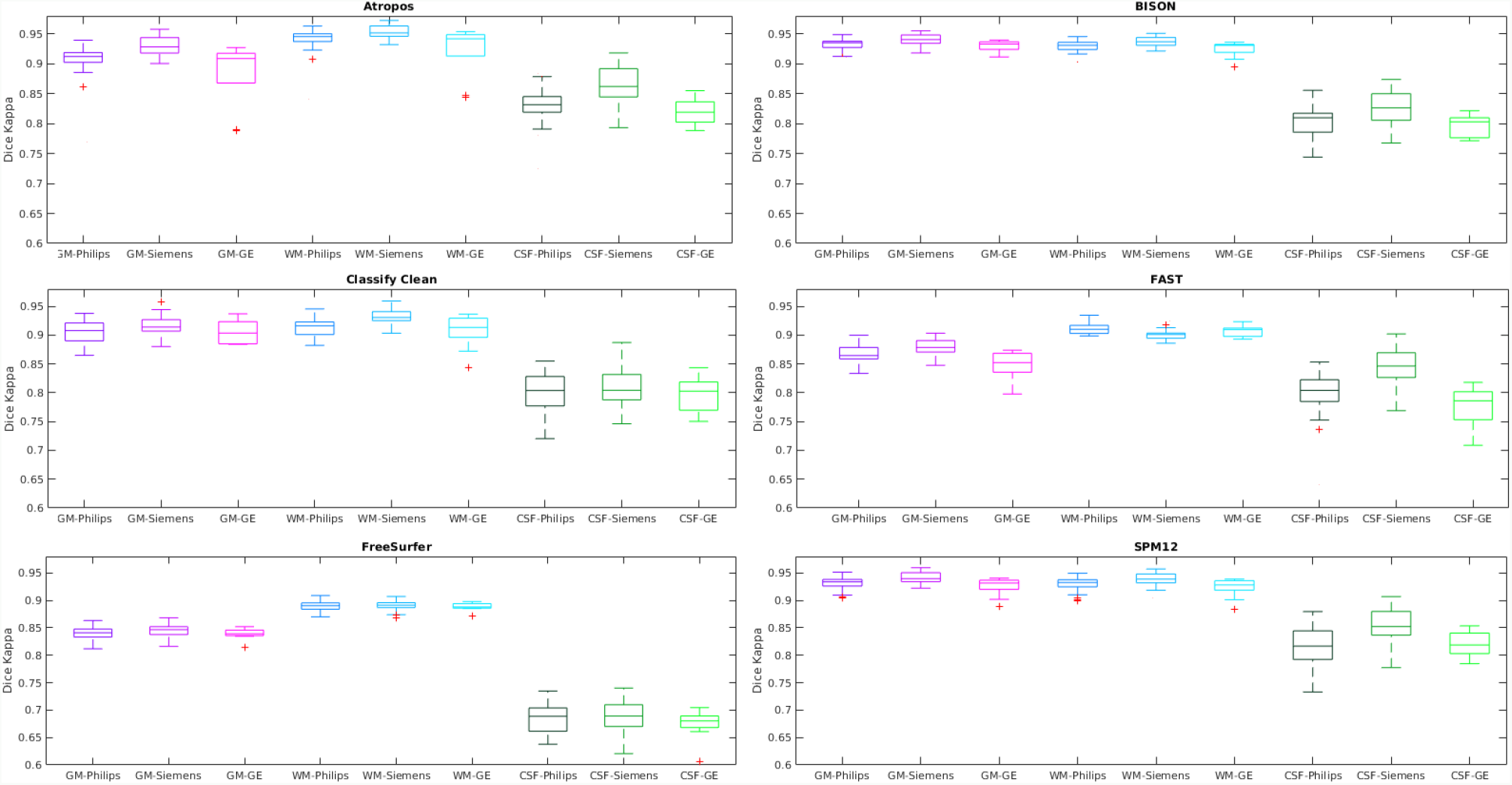
Representation of Dice Kappa results from Table 1 for individual image segmentation, when compared to the silver standard segmentation mask, for tissue volumes across scanner manufacturers. GM=Gray Matter. WM=White Matter. CSF= CerebroSpinal Fluid.

Across tissue types, BISON had the lowest overall variability in Dice Kappas, followed by SPM12. For GM, BISON had the highest overall Dice Kappa (0.94±0.01), followed by SPM12 and Atropos; while Atropos had the highest overall Dice Kappa for WM (0.94±0.02), followed by BISON and SPM12. To assess the statistical significance of the results, paired *t*-tests were performed on the Dice Kappa values of all pairs of segmentation techniques, and the resulting p-values were corrected for multiple comparisons (FDR). Figure 4 shows the negative logarithm of the FDR corrected p-values. All methods tended to have higher Dice Kappa values for scans from Siemens compared to Philips and GE.

**Figure 4.**
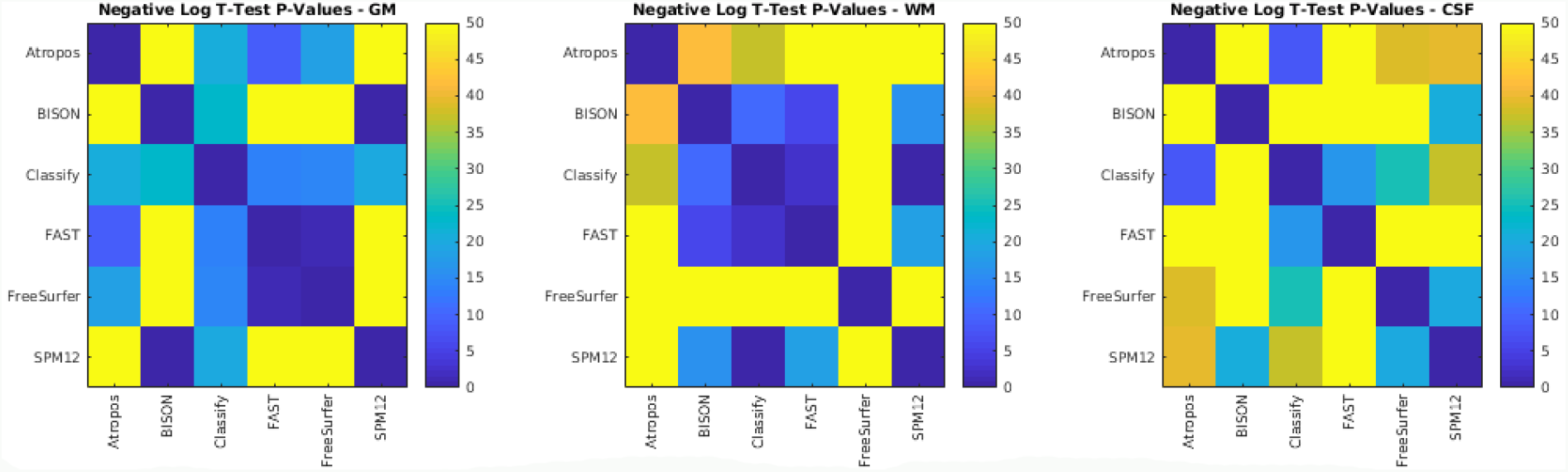
Negative logarithm of FDR corrected p-values of paired t-tests between Dice Kappa values of segmentation technique pairs. Values higher than 1.3 are statistically significant.

Table 2 shows average estimated volumes for GM, WM, and CSF from each scanner, as well as the averages separately for each manufacturer. Table 3 shows the estimated volumes differences across manufacturers, obtained from the linear regression model (eq. 1). Significant differences (after FDR correction) are displayed in bold font. Figure 5 shows boxplots of the acquired volumes for each tissue type across scanner manufacturers. Figure S.3 shows boxplots of the acquired volumes across scanner models.

**Table 2.** Average tissue volumes for each method. Values indicate mean ± standard deviation normalized by the intracranial volume.

| Tissue | Gray Matter |  |  |  | White Matter |  |  |  | Cerebrospinal Fluid |  |  |  |
| --- | --- | --- | --- | --- | --- | --- | --- | --- | --- | --- | --- | --- |
| Method | Overall | GE | Philips | SIEMENS | Overall | GE | Philips | SIEMENS | Overall | GE | Philips | SIEMENS |
| Atropos | 47.64±1.9 | 46.25±4.9 | 46.97±1.2 | 48.24±0.7 | 39.97±1.8 | 41.04±5.3 | 40.05±1.3 | 39.80±0.7 | 12.39±0.9 | 12.70±0.7 | 12.98±0.8 | 11.96±0.7 |
| BISON | 57.60±0.4 | 58.25±0.3 | 57.38±0.3 | 57.67±0.5 | 34.49±0.4 | 33.72±0.2 | 34.57±0.4 | 34.55±0.4 | 7.90±0.3 | 8.01±0.2 | 8.05±0.3 | 7.78±0.3 |
| Classify | 53.11±3.0 | 54.93±3.8 | 53.12±3.8 | 52.91±2.4 | 33.05±1.7 | 32.77±1.9 | 33.71±2.1 | 32.69±1.3 | 13.83±2.0 | 12.30±2.2 | 13.16±2.2 | 14.32±1.7 |
| FAST | 49.35±2.2 | 47.55±1.8 | 47.92±2.4 | 50.45±1.3 | 33.74±1.2 | 34.09±0.8 | 34.56±1.4 | 33.31±1.1 | 16.90±1.8 | 18.35±1.9 | 17.50±2.0 | 16.24±1.3 |
| FreeSurfer | 49.80±0.7 | 49.89±0.5 | 49.24±0.6 | 50.13±0.6 | 39.52±0.8 | 39.39±1.2 | 39.92±0.9 | 39.33±0.5 | 10.67±0.9 | 10.71±1.2 | 10.83±0.9 | 10.53±0.8 |
| SPM12 | 57.76±1.3 | 57.21±2.5 | 57.95±1.5 | 57.73±0.8 | 33.08±1.1 | 33.73±2.0 | 33.57±1.3 | 32.72±0.6 | 9.12±0.9 | 9.03±1.1 | 8.48±0.7 | 9.50±0.6 |

**Table 3.** Volume differences across scanner manufacturers. Values represent beta estimate. Significant results after FDR correction (threshold =0.05) are shown in bold font. GM=Gray Matter. WM=White Matter. CSF= CerebroSpinal Fluid. FDR= False Discovery Rate.

| Method | Gray Matter |  |  | White Matter |  |  | Cerebrospinal Fluid |  |  |
| --- | --- | --- | --- | --- | --- | --- | --- | --- | --- |
| t-stat | GE vs. Philips | GE vs. SIEMENS | SIEMENS vs. Philips | GE vs. Philips | GE vs. SIEMENS | SIEMENS vs. Philips | GE vs. Philips | GE vs. SIEMENS | SIEMENS vs. Philips |
| Atropos | -0.72 | **-2.05** | **1.33** | 0.99 | 1.32 | -0.32 | -0.28 | **0.73** | **-1.01** |
| BISON | **0.88** | **0.64** | 0.23 | **-0.84** | **-0.87** | 0.03 | -0.03 | 0.23 | -0.26 |
| Classify | 1.81 | 2.15 | -1.02 | -0.95 | 0.07 | -1.02 | -0.86 | **-2.15** | **1.34** |
| FAST | -0.92 | **-3.54** | **2.63** | -0.36 | **1.03** | **-1.39** | 1.28 | **2.51** | -1.24 |
| FreeSurfer | 0.65 | -0.23 | **0.88** | 0.58 | -0.53 | **1.11** | -0.12 | 0.14 | -0.26 |
| SPM12 | -0.74 | -0.54 | -0.21 | 0.16 | **1.07** | **-0.91** | 0.58 | -0.53 | **1.11** |

**Figure 5.**
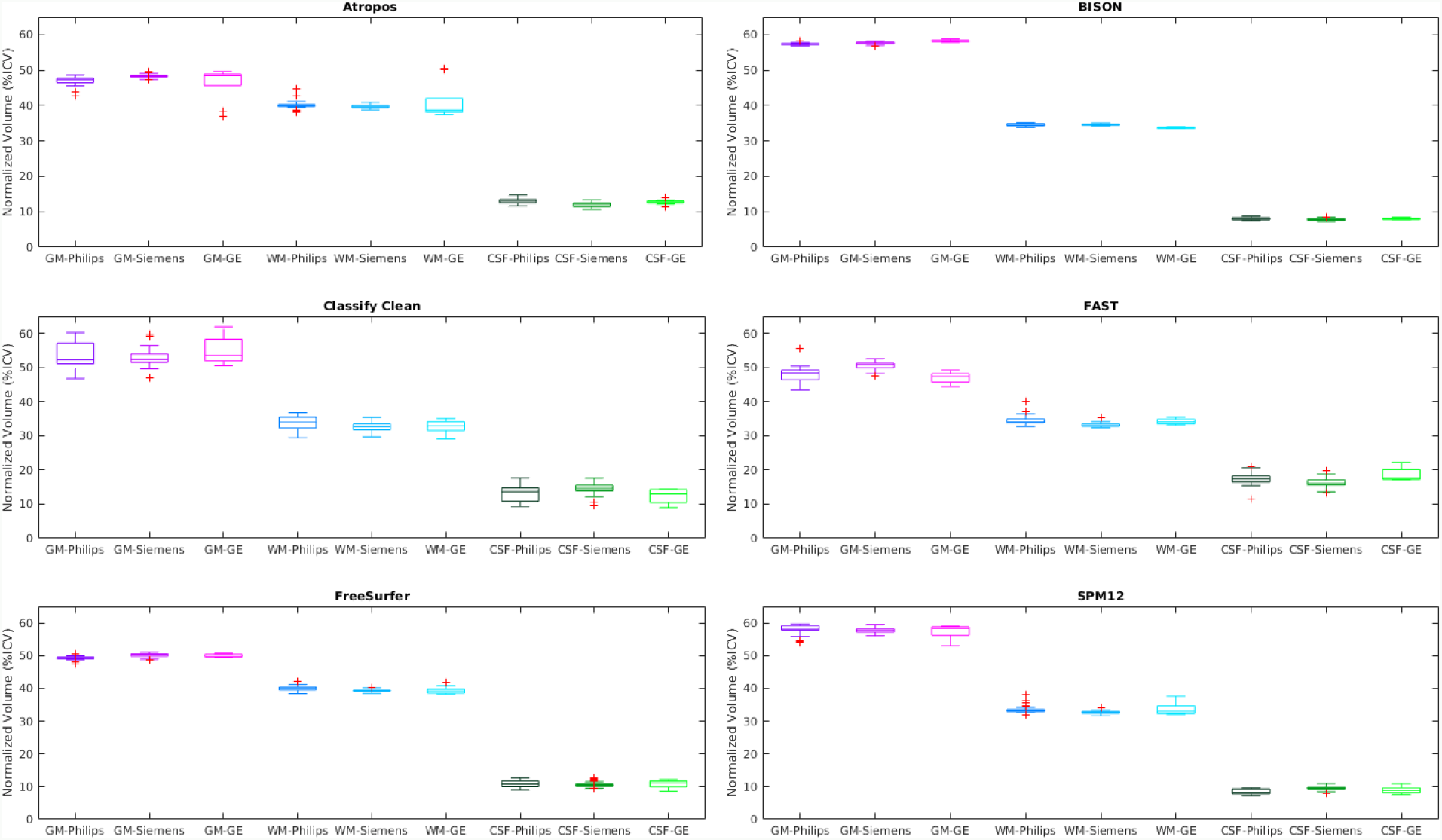
Tissue volumes across scanner manufacturers. GM=Gray Matter. WM=White Matter. CSF= CerebroSpinal Fluid.

BISON had the lowest overall variability, followed by FreeSurfer, and SPM12 (Table 2). Atropos estimated significantly greater GM volumes and significantly lower CSF volumes in Siemens scans, compared with the other two manufacturers (Table 3). BISON estimated significantly greater GM volumes and significantly lower WM volumes in GE scans, compared with the other two manufacturers. FAST estimated significantly lower GM volumes and significantly higher CSF volumes in GE scans compared with Siemens, and significantly greater GM volumes and significantly lower WM volumes in Siemens scans compared with Philips. Similarly, FreeSurfer estimated significantly greater GM volumes and significantly lower WM volumes in Siemens scans compared with Philips. SPM12 estimated significantly lower WM volumes and higher CSF volumes in Siemens scans compared with Philips.

Table 4 shows the estimated differences between the segmented tissue volumes by each method for scans in low, medium, and high tertiles of SNR, obtained from the linear regression model (eq. 2). Significant differences (after FDR correction) are displayed in bold font. Except for FreeSurfer, all methods had significantly different CSF estimates across high and low SNR tertiles. FAST and FreeSurfer had significantly higher WM volume estimates for high versus medium and low tertiles as well as corresponding significantly lower GM volume estimates. SPM12 had significantly higher WM volume estimates for high versus medium and low tertiles as well as corresponding significantly lower CSF volume estimates. Atropos had a significantly lower estimate for GM volume of high versus low tertiles, and a corresponding higher estimate for WM volume.

**Table 4.**
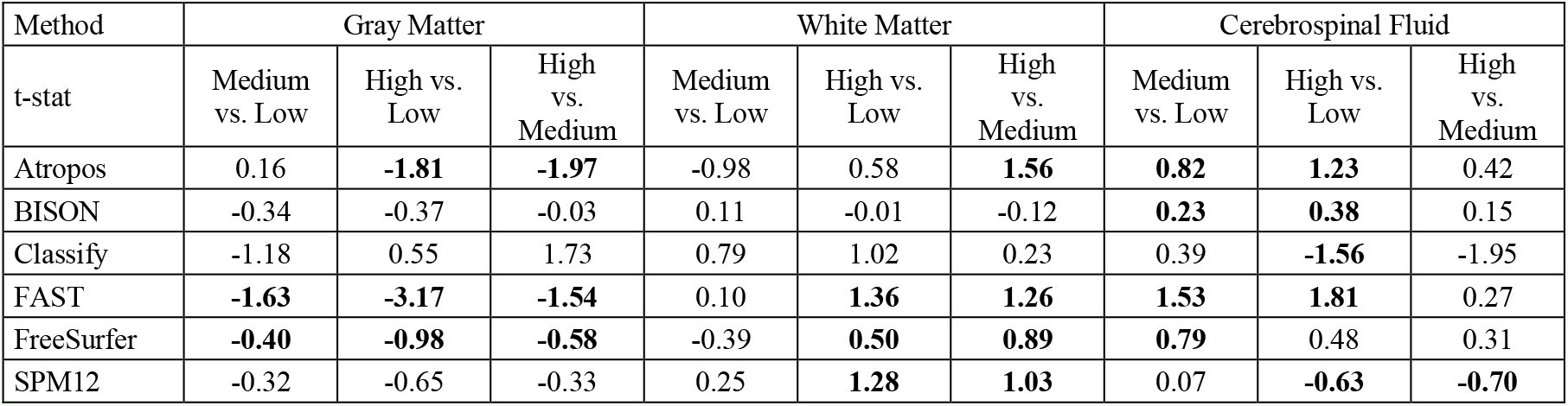
Impact of SNR on estimated volumes. Values represent beta estimate. Significant results after FDR correction (threshold =0.05) are shown in bold font. SNR= Signal to Noise Ratio. GM=Gray Matter. WM=White Matter. CSF= CerebroSpinal Fluid. FDR= False Discovery Rate.

Using the volume estimates from Table 2, the sample sizes necessary to detect a 1% reduction in the tissue volumes were estimated (Table 5, 80% power, 2-tailed significance). BISON had the smallest sample size requirement across all scanners and tissue types, followed by FreeSurfer, and SPM12. As expected, the necessary sample size decreased for all methods and tissue types when using data from one specific scanner.

**Table 5.** Estimated sample size to detect a 1% reduction in the tissue volumes (80% power, 2-tailed significance).

| Tissue | Gray Matter |  |  |  | White Matter |  |  |  | Cerebrospinal Fluid |  |  |  |
| --- | --- | --- | --- | --- | --- | --- | --- | --- | --- | --- | --- | --- |
| Method | Overall | GE | Philips | SIEMENS | Overall | GE | Philips | SIEMENS | Overall | GE | Philips | SIEMENS |
| Atropos | 127 | 93 | 21 | 12 | 162 | 136 | 32 | 16 | 417 | 27 | 107 | 115 |
| BISON | 6 | 3 | 4 | 6 | 13 | 3 | 6 | 9 | 116 | 8 | 41 | 69 |
| Classify | 253 | 41 | 144 | 94 | 210 | 29 | 110 | 73 | 1644 | 259 | 775 | 631 |
| FAST | 158 | 11 | 72 | 32 | 102 | 7 | 48 | 51 | 893 | 89 | 364 | 288 |
| FreeSurfer | 18 | 4 | 7 | 9 | 35 | 10 | 17 | 10 | 561 | 103 | 193 | 260 |
| SPM12 | 42 | 18 | 21 | 11 | 89 | 31 | 44 | 18 | 767 | 122 | 191 | 180 |

## 4. Discussion

In this paper, we assessed the variability in tissue segmentation results for six publicly available and widely used tissue classification methods in the context of a large body of images for a single volunteer acquired on multiple scanner manufacturers and models across time. Such assessments are particularly important for the field of neuroimaging, given at present many researchers are transitioning to using large multi-center and multi-scanner databases in order to test their hypotheses with sufficient statistical power, and/or using machine learning techniques that require a large array of data. Our comparisons provide a benchmark for the expected variability and systematic differences between results obtained from the same image processing pipeline for scans from different centers, as well as differences that can be expected when comparing volumetric results obtained from different pipelines. If such differences are systematic and consistent, one can select the algorithm with lowest variability, or adjust for the differences when using data from multiple scanner manufacturers.

BISON GM segmentations showed the highest overlap (i.e. average Dice Kappa) with the silver standard obtained based on the average template, followed by SPM12 and Atropos. Atropos had the highest overlap for the WM segmentations, followed by BISON and SPM12. The overlap comparisons in general tended to agree with the volumetric comparisons, except for FreeSurfer results. This can be explained by the fact that the average template was preprocessed (i.e. denoised, non-uniformity corrected, and intensity normalized) prior to the FreeSurfer processing, whereas the individual scans processed by FreeSurfer were not. However, since FreeSurfer produced significantly poorer results when segmenting previously preprocessed images, we were not able to compare FreeSurfer results on scans that were consistently preprocessed.

BISON had the lowest variability in its estimated tissue volumes, followed by FreeSurfer and SPM12 (Table 2). This lower variability might be due to the fact that BISON itself was trained based on a multi-center and multi-scanner dataset, and therefore was able to deal with some of the variabilities across scanners.

SNR had a significant impact on many of the estimated volumes. All methods had significantly different CSF estimates across high and low SNR tertiles. FAST, FreeSurfer, and SPM12 also had significantly higher WM volume estimates for high versus medium and low tertiles as well as corresponding significantly lower GM volume estimates. Overall, BISON had the smallest amount of difference between the volumes estimated across SNR tertiles, followed by FreeSurfer. This is an important concern, particularly when using data from older 1.5T scanners which tend to have lower SNRs. These results also signify the importance of acquiring data with the best possible SNR to minimize the consequent variability in volumetric measurements.

BISON had the smallest sample size requirement across all scanners and tissue types, followed by FreeSurfer, and SPM12. As expected, the necessary sample size decreased for all methods and tissue types when using data from one specific scanner. This is an important concern when designing multi-scanner studies that acquire data using scanners from different manufacturers.

One of the limitations of this study is the inconsistent distribution of the data across different scanners. Out of the 90 scans used in this study, nine scans were acquired on GE, 31 on Philips, and 50 on Siemens. In addition, mean ± standard deviation for age was 45.49 ± 1.07 for GE scans, 42.28 ± 2.85 for Philips scans, and 44.85 ± 1.26 for Siemens scans. The differences between age at scan for Philips with the other two manufacturers were statistically significant (p<0.002). These differences might introduce some variability into the scanner comparisons results that are not caused by scanner differences.

In this paper, we have compared the performance of six publicly available, widely used tissue classification methods on a travelling human phantom dataset, containing 90 scans across 28 sites, and with 12 different scanner models. Our comparisons provide a practical benchmark on the reliability of each technique, and the amount of variability that can be expected across scanners from different manufacturers and SNR levels when using multi-center and multi-scanner datasets.

## Acknowledgements

Part of the data used in this article were obtained from the Consortium pour l’identification précoce de la maladie Alzheimer - Québec (CIMA-Q; cima-q.ca) and from the Canadian Consortium on Neurodegeneration in Aging (CCNA; www.ccna-ccnv.ca). As such, the investigators within the CIMA-Q contributed to the design, the implementation, the acquisition of clinical, cognitive, and neuroimaging data and biological samples. A list of the CIMA-Q investigators is available on www.cima-q.ca. CIMA-Q is financed through the Fonds de recherche du Québec – Santé/Pfizer Canada Innovation Fund (#27239). The CCNA is financed through the Canadian Institutes for Health Research (2014–2019) with funding from several partners. In addition to CIMA-Q and CCNA, other organizations and projects have contributed to the elaboration of the CDIP protocol, namely the ONDRI (ondri.ca) and courtesy scans at MR manufacturers. The Ontario Brain Institute is financed by the Government of Ontario and the Ontario Brain Institute Foundation. Financial support for S.D. for travel in order to obtain scans was obtained from the Canadian Institutes for Health Research (#117121), and the Fonds de recherche du Québec – Santé/Pfizer Canada - Pfizer-FRQS Innovation Fund (#25262). We also wish to thank the Cuban Neuroscience Center, specifically its Human Brain Mapping Unit, and the University of Electronic Science and Technology of China for their interest in importing the Canadian Dementia Imaging Protocol, with support from the Fonds de recherche du Québec – Santé tri-national Neuroinformatics and Neuroimaging collaboration program.

## Supplementary Materials

**Figure S.1.**
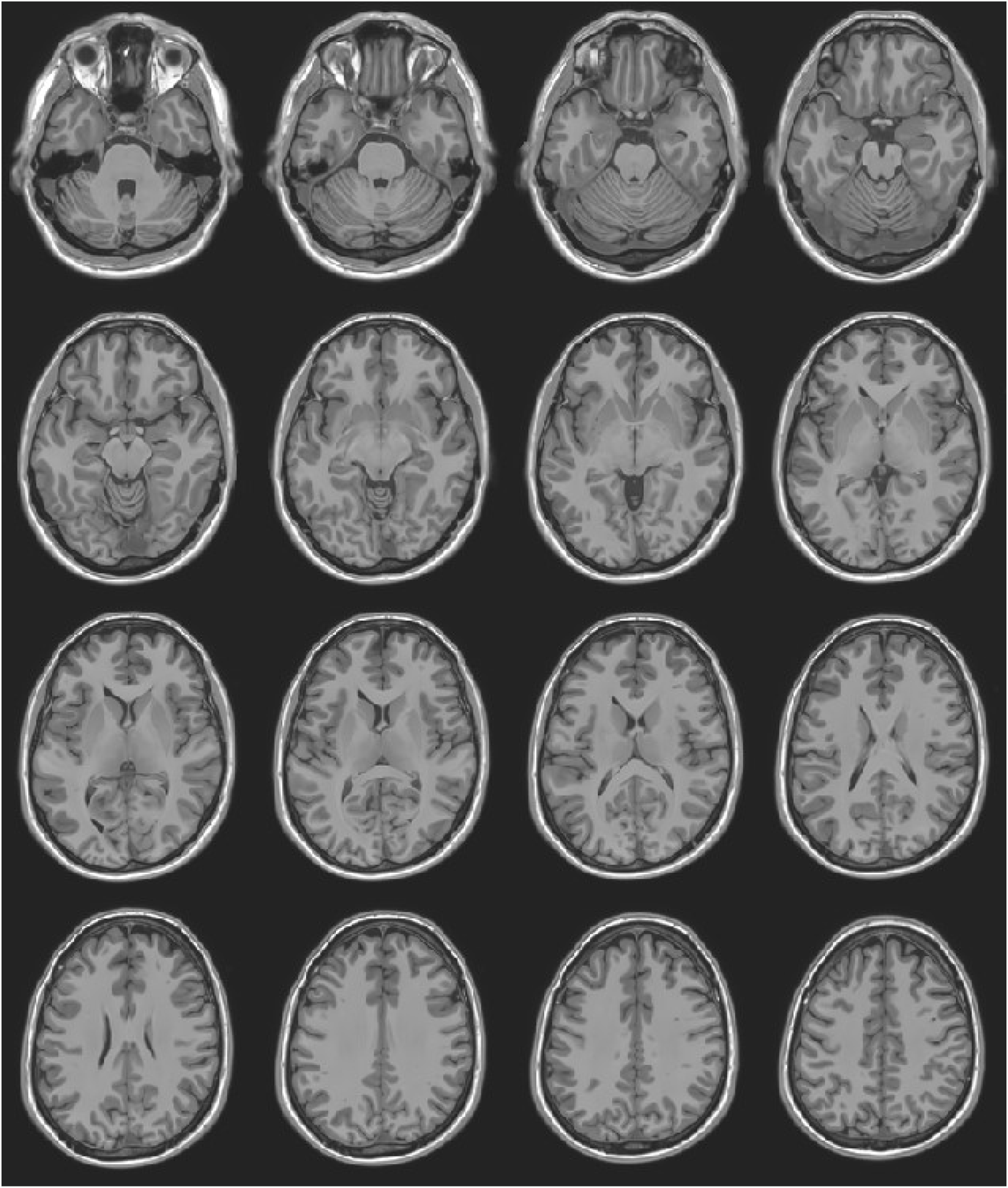
Axial slices showing the average SIMON brain.

**Figure S.2.**
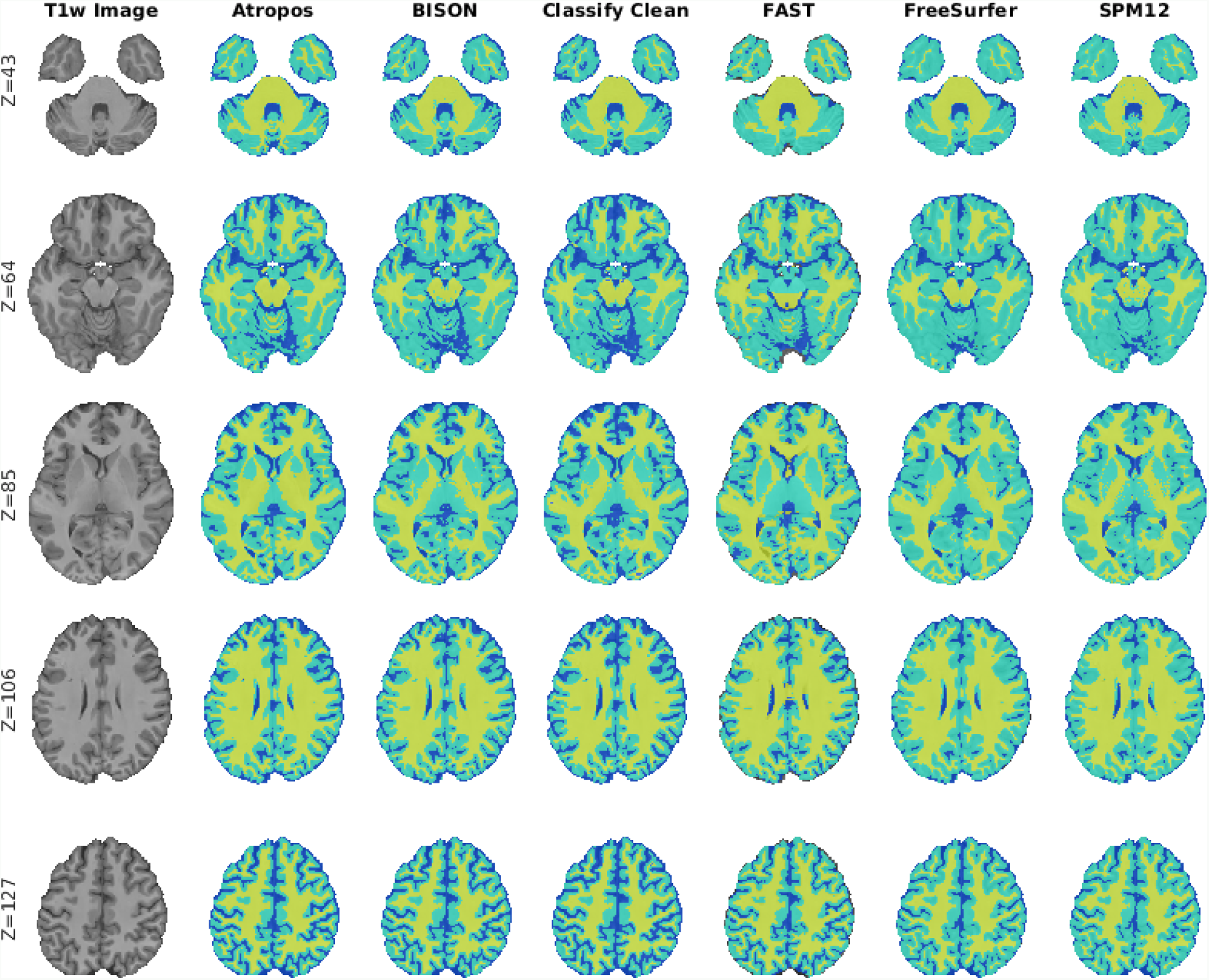
Axial slices comparing segmentations from Atropos, BISON, Classify_Clean, FAST, FreeSurfer, and SPM12 for one scan (Philips Intera 3T scanner).

**Figure S.3.**
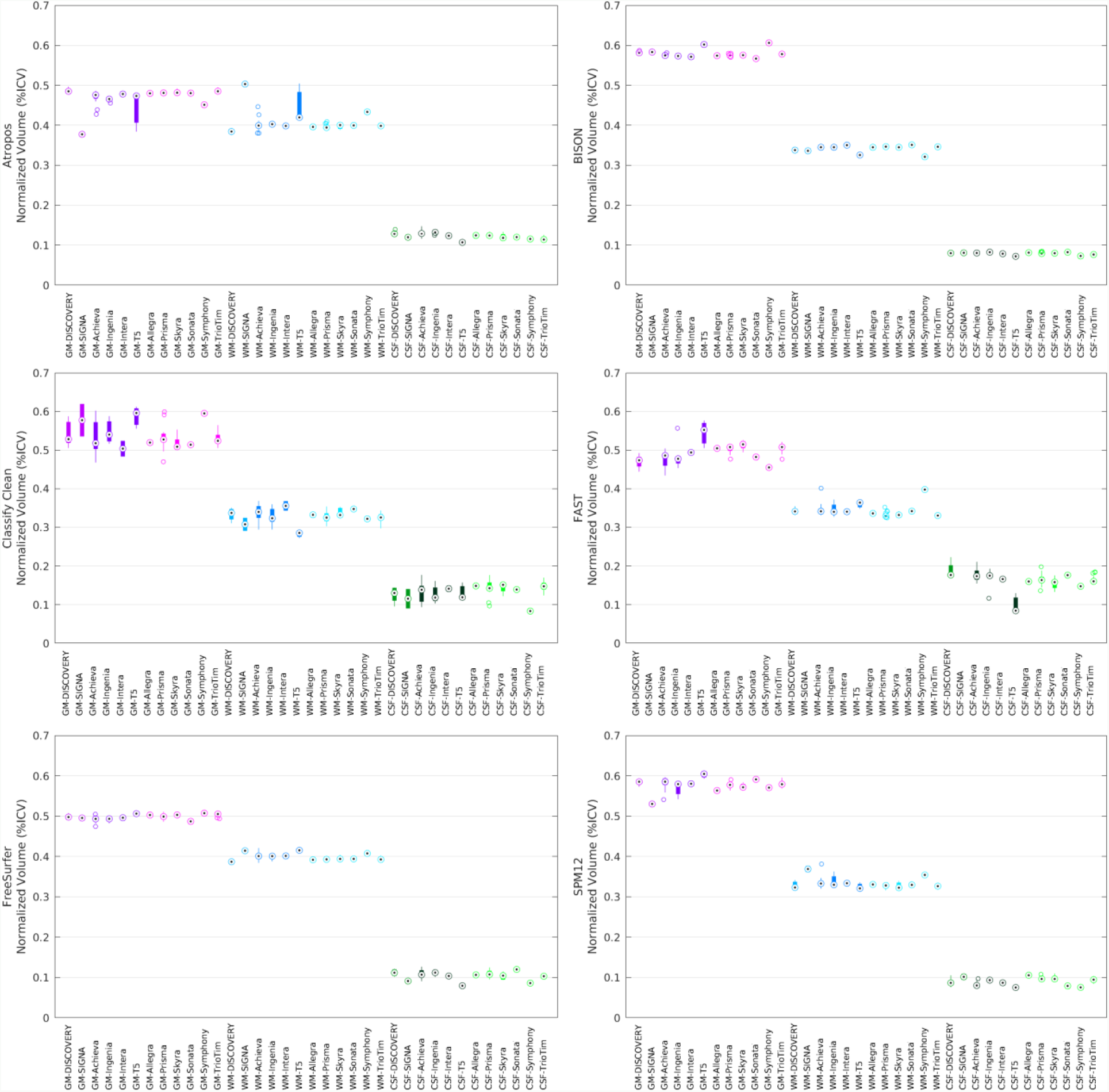
Estimated tissue volumes across scanner models. GM=Gray Matter. WM=White Matter. CSF= CerebroSpinal Fluid.

